# Gene expression analysis suggests immunosuppressive roles of endolysosomes in glioblastoma

**DOI:** 10.1101/2023.08.31.555740

**Authors:** Michael A Sun, Haipei Yao, Qing Yang, Christopher J. Pirozzi, Vidyalakshmi Chandramohan, David M. Ashley, Yiping He

**Affiliations:** The Preston Robert Tisch Brain Tumor Center, Duke University Medical Center, Durham, NC 27710, USA; Department of Pathology, Duke University Medical Center, Durham, NC 27710, USA; Pathology Graduate Program, Duke University Medical Center, Durham, NC 27710, USA; Duke University School of Nursing, Durham, NC 27710, USA; Department of Neurosurgery, Duke University Medical Center, Durham, NC, 27710 USA

**Keywords:** glioblastoma, lysosome, endolysosome, tumor-associated macrophage

## Abstract

Targeting endolysosomes is a strategy extensively pursued for treating cancers, including glioblastomas (GBMs), mainly on the basis that the intact function of these subcellular organelles is key to tumor cell autophagy and survival. Here, by gene expression analyses and cell type abundance estimation in GBMs, we showed that genes associated with the endolysosomal machinery are more prominently featured in non-tumor cells in GBMs than in tumor cells, and that tumor-associated macrophages represent the primary immune cell type that contributes to this trend. Further analyses found an enrichment of endolysosomal pathway genes in immunosuppressive (pro-tumorigenic) macrophages, such as M2-like macrophages or those associated with worse prognosis in glioma patients, but not in those linked to inflammation (anti-tumorigenic). Specifically, genes critical to the hydrolysis function of endolysosomes, including progranulin and cathepsins, were among the most positively correlated with immunosuppressive macrophages, and elevated expression of these genes is associated with worse patient survival in GBMs. Together, these results implicate the hydrolysis function of endolysosomes in shaping the immunosuppressive microenvironment of GBM. We propose that targeting endolysosomes, in addition to its detrimental effects on tumor cells, can be leveraged for modulating immunosuppression to render GBMs more amendable to immunotherapies.

## Introduction

Lysosomes are hydrolase-rich intracellular organelles critical for normal development and cell physiology in mammalian cells^1^. When fused with intracellular membrane-enclosed organelles, such as autophagosomes and endosomes, the fused compartments (i.e., autophagolysosomes and endolysosomes) provide the acidic environment in which hydrolases (e.g., cathepsin proteases^2^) become enzymatically active and break down lipids and proteins to recycle these macromolecules and maintain their homeostasis. The endolysosomal system has also been shown to be essential for cancer cell proliferation, survival, and autophagy^3,4^, and thus has been extensively investigated as a therapeutic target for cancer treatments. Various strategies for disrupting endolysosomal functions have been developed, including inducing lysosome membrane permeabilization (LMP; caused by cationic amphiphilic drugs), targeting autophagy (e.g., chloroquine and its derivative hydroxychloroquine), and inhibiting cathepsins and vacuolar H^+^-ATPases, which are acidic pH-maintaining lysosome membrane proteins^3,4^. Such strategies were developed primarily based on consequences from cancer cell killing, and their efficacy is frequently assayed centering on cancer cells, leaving their direct effect on non-tumor cells and/or the immune microenvironment unclear^3-6^.

## Methods

The mRNA-seq data (gene expression in transcripts per million (tpm)) from The Cancer Genome Atlas GBM (TCGA-GBM) dataset were obtained from the GDC data portal (https://portal.gdc.cancer.gov/). Cell type deconvolution was performed using EPIC (Estimating the Proportions of Immune and Cancer cells, via http://epic.gfellerlab.org)^7^, and quanTIseq^8^, xCELL^9^ and CIBERSORT (absolute mode)^10^, which were run from the web interface TIMER 2.0 (http://timer.cistrome.org/)^11^. Pathway enrichment analysis was performed via ShinyGO (version 0.77: http://ge-lab.org/go/)^12^ using the Gene Ontology - Cellular Component database, and validated by independent pathway enrichment analysis performed via g:Profiler^13^. For the analysis of TCGA GBM samples, all protein-coding genes identified in the mRNA-seq data were used as the background gene set, and a false discovery rate (FDR) cutoff of 0.05 was used to define significantly enriched pathways. Correlations between immune genes and endolysosomal machinery genes were performed using gene expression tpm or obtained from the glioma dataset visualization web application Gliovis (http://gliovis.bioinfo.cnio.es/)^14^. For the pathway enrichment analysis of *SIGLEC9*^+^ TAMs, differentially expressed genes (FDR <= 0.05) in *SIGLEC9*^+^ versus *SIGLEC9*^-^ TAMs were obtained from a previous study^15^, and all known protein-coding genes were used as the background gene set for the pathway enrichment analysis. The association of gene expression with patient survival of patients from CGGA (Chinese Glioma Genome Atlas^16^) and TCGA-GBM datasets were obtained from Gliovis^14^. Heatmaps were generated using Heatmapper (http://www.heatmapper.ca; clustering method: single linkage; distance measurement method: Euclidean)^17^. For all statistical analyses for gene expression correlations, p-values were determined using GraphPad and p < 0.05 was considered statistically significant.

## Results

We initially sought to pinpoint the most prominent cellular components and processes in glioblastoma (GBM), one of the most lethal cancers, to facilitate the identification of therapeutic targets. We utilized the mRNA-seq data of 162 GBM patients from the TCGA-GBM dataset, and estimated the fractions of tumor cells and various types of immune cells using the EPIC (Estimating the Proportions of Immune and Cancer cells) method^7^. We then correlated cancer cells and the remaining cells (i.e., immune cells, endothelial cells, and cancer-associated fibroblasts (CAFs)) with the expression level of each gene (total of 19961 protein-coding genes were examined). Genes that were most positively correlated with cancer cells (cutoff used: r >= 0.3, p <= 0.0001; n = 1257 genes) were subjected to pathway enrichment analyses using the Gene Ontology-Cellular Component (GO-CC) database. The results indicated that, as expected, the most prominent cellular components in cancer cells included those involved in DNA replication (e.g., replisome), gene transcription (e.g., RNA polymerase core complex), protein translation (e.g., tRNA synthetase complex and small ribosomal subunits), and epigenetic reprogramming (e.g., histone acetyltransferase complex and methylosome), all of which were cellular machinery essential to the malignant properties of tumor cells (**Fig. 1A**). Interestingly, similar analysis of genes most negatively correlated with cancer cells (r <= -0.5, p < 0.00001; n = 654) identified an entirely different list of cellular components, including those associated with tumor microenvironment (e.g., protein complex of cell adhesion and collagen trimers), granules and granule membranes, and most notably, primary lysosomes and endolysosomes (**Fig. 1B**). These results were corroborated by a subsequent analysis using another deconvolution method, quanTIseq^8^ (**Supplementary Fig. 1A-B**), suggesting that the endolysosome system not only represents a promising therapeutic target against cancer cells, but might also play significant roles in orchestrating non-tumor cells in GBMs.

**Fig. 1.**
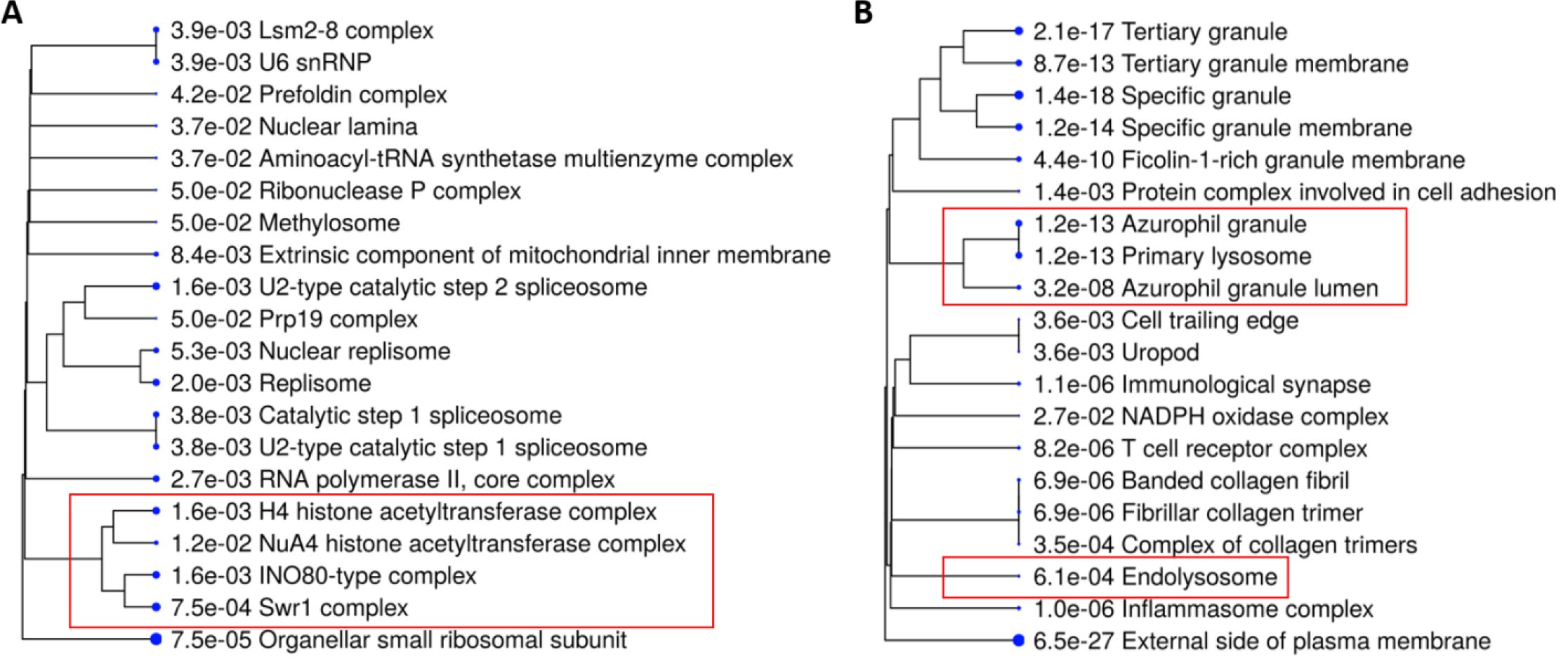
Gene expression analysis suggests tumor and non-tumor cells in GBMs featured distinct cellular components. The mRNA-seq data (protein-coding genes, tpm) of 162 TCGA GBM samples were subjected to EPIC analysis for estimating fractions of each type of cells. **(A)** GO-Cellular Component enrichment analysis of genes positively correlated with cancer cells (annotated as “Other Cells”) (r >= 0.3; n = 1257). **(B)** GO-Cellular Component enrichment analysis of genes negatively correlated with cancer cells (r <= -0.5; n = 654). (A-B) Dendrograms show the top 20 cellular components sorted by fold enrichment, and the size of each dot denotes the relative number of genes in each cellular component.

Among non-tumor cells identified in the deconvolution analysis, three types of cells were among the most abundant: macrophages, endothelial cells, and CAFs (**Supplementary Fig. 2A**). To assess which of these three cell types contributed to the above enrichment of lysosomes/endolysosomes, we further performed GO-Cellular Component enrichment analysis for genes positively correlated with each cell type. We found that genes most positively correlated with macrophages (r >= 0.5, p < 0.00001; n = 543) were enriched for lysosomes/endolysosomes (**Fig. 2A**), and these cellular components were similarly identified when only the genes with the strongest positive correlation (r >= 0.7, p < 0.00001; n = 208) were included for pathway enrichment analysis (**Fig. 2B**). In contrast, similar analyses applied to endothelial cells and CAFs did not identify endolysosomal compartments, while cellular components characteristic of each cell type, such as apical plasma membrane and collagen-containing extracellular matrix, were respectively enriched as expected (**Supplementary Fig. 2B-C**). Finally, the enrichment of lysosomes/endolysosomes in macrophages was further validated by independent estimation of cell type abundance using another deconvolution method (xCELL)^9^ and subsequent pathway analysis (**Supplementary Fig. 2D**). Collectively, these results suggest that lysosomes/endolysosomes are prominent cellular components of GBM tumor-associated macrophages (TAMs), a major, heterogeneous population of cells that are key to shaping immune microenvironment and mediating immunosuppression in GBM^18-20^.

**Fig. 2.**
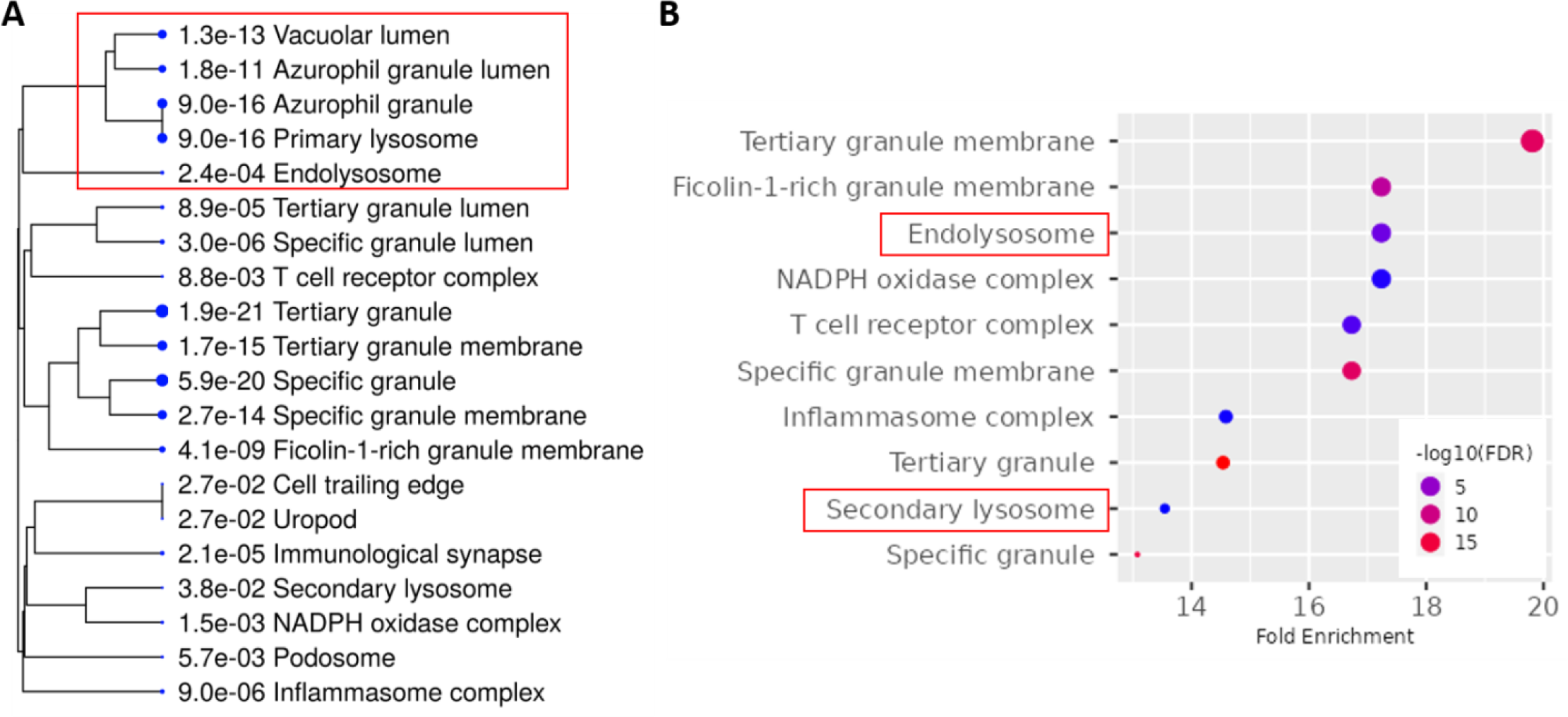
Tumor-associated macrophages are the major cell type that contributes to the prominent feature of endolysosomal machinery in GBMs. **(A-B)** The mRNA-seq data (protein-coding genes, tpm) of 162 GBM samples were subjected to EPIC analysis for estimating fractions of each type of cell. **(A)** GO-Cellular Component enrichment analysis of genes positively correlated with macrophages (r >= 0.5; n = 543). The dendrogram shows the top 20 cellular components sorted by fold enrichment, and the size of each dot denotes the relative number of genes in each cellular component. **(B)** GO-Cellular Component enrichment analysis of genes identified as displaying strong positive correlations (r >= 0.7; n = 208) with macrophages. The chart plot shows the top 10 enriched cellular components (FDR cutoff = 0.05, sorted by fold enrichment). The size of each dot corresponds to the fold enrichment for each cellular component as shown on the x-axis.

The heterogeneity of TAMs led us to ask whether the feature of lysosomes/endolysosomes is uniformly true in all TAMs, or else more prominent in specific subsets of TAMs. Answering this question is important, as it can provide insight into not only the pathological roles, mechanism of action, and regulation of various subtypes of TAMs in GBMs, but also the future therapeutic implications. For example, if the endolysosomal system is more uniquely associated with the anti-tumorigenic TAMs (“good” TAMs), undesired outcomes may be inflicted when therapeutically disrupting the function of these organelles. In contrast, if they are characteristic of and essential to pro-tumorigenic (i.e., immunosuppressive) TAMs, then targeting lysosomes/endolysosomes can likely mitigate these “bad” cells, providing a beneficial effect in addition to direct killing of tumor cells. To address this question, we first used CIBERSORT to estimate the abundance of two major types of macrophages, M1-like and M2-like macrophages, representing the more anti-tumorigenic and the more pro-tumorigenic TAMs, respectively^19^. We performed similar cellular component enrichment analysis for genes positively correlated with the fraction of each type of TAMs. This analysis revealed that genes positively correlated with M2 macrophages were enriched with components of lysosomes/endolysosomes (**Supplementary Fig. 3A**). Intriguingly, the cellular components enriched in genes positively correlated with M1 macrophages were dominated by those of cell-cell interactions and cell-extracellular communication (e.g., cell surface proteins and protein complexes involved in cell adhesion) (**Supplementary Fig. 3B**), suggesting that various types of macrophages likely rely on distinct cellular organelles and compartments to exert their distinct roles in GBMs, such as immunosuppression versus anti-tumor response.

While subtyping TAMs using the M1/M2 paradigm is a useful means of illustrating their opposite roles in cancers, it has emerged that the *in vivo* evidence supporting this dichotomous classification of TAMs remains absent in GBM, and that these cells are more heterogenous than previously thought^18^. Most notably, a recent single-cell RNA sequencing study has defined nine myeloid clusters (MC) in human gliomas (including low-grade glioma, newly diagnosed GBMs, and recurrent GBMs), and identified those that were correlated with better survival (e.g., MC2 and MC7, microglia clusters proposed to be anti-tumorigenic) or worse survival (e.g., MC3 and MC5, macrophage clusters proposed to be immunosuppressive and pro-tumorigenic) in GBM patients^21^. We performed similar analysis for marker genes for these two groups of myeloid cells, and found that lysosomes/endolysosomes were enriched only in the pro-tumorigenic myeloid clusters (**Fig. 3A-B**). Further individual examination of markers genes for MC3 (n = 55) and MC5 (n = 254) found that the enrichment of the lysosome/endolysosome components, while observed in both MCs, was particularly dominant in MC3 (**Supplementary Fig. 3C**), the cluster that was more strongly associated with poor survival in GBM patients^21^. In all, 21 (38%) of the 55 marker genes for MC3 were defined as being explicitly associated with GO-Cellular Components of lysosomes and/or endosomes (GO:0005764, GO:0005768, and GO:0043202), and 16 (29%) of them encode proteins that have been determined to reside in endosomes and/or lysosomes per the COMPARTMENTS database^22^, including proteins directly involved in the degradation process in endolysosomes, such as hydrolases (e.g., *CTSD, CTSL, CTSZ*, and *FUCA1*) and progranulin (*GRN*), a non-enzymatic protein recently found to be indispensable for lysosomal hydrolysis^23,24^ (**Fig. 3C** and **Supplementary Table 1**). In-depth examination of the MC3 genes further corroborates the roles of endolysosomes in GBM pathogenesis and, specifically, in immunosuppression. First, we assessed the relationship between the 21 MC3 genes and a set of four genes representative of GBM’s immune suppressive microenvironment: the immunosuppressive TAM markers *MRC1* (CD206) and *CD163*^18^ and the immunosuppressive cytokines *IL10* and *TGFB1*^18,20^, and found overwhelmingly positive correlations between these two groups of genes (**Fig. 4A-B, Supplementary Fig. 4**, and **Supplementary Table 1**). Subsequently, we examined the association of each of the 21 MC3 genes with GBM patient’s survival from the TCGA and CGGA datasets, and found that high expression level of each of the 11 genes was significantly associated with worse overall survival in at least one of the datasets (**Supplementary Table 1**). Among them, six genes were found to be significantly linked to worse survival in the TCGA dataset while displaying similarly significant association or at least a consistent trend in the CGGA dataset, including five genes that encode proteins that primarily localized in endolysosomes (*CTSD, CTSZ, FUCA1, GRN*, and *IFI30*) (**Fig. 4C-D** and **Supplementary Table 1**).

**Fig. 3.**
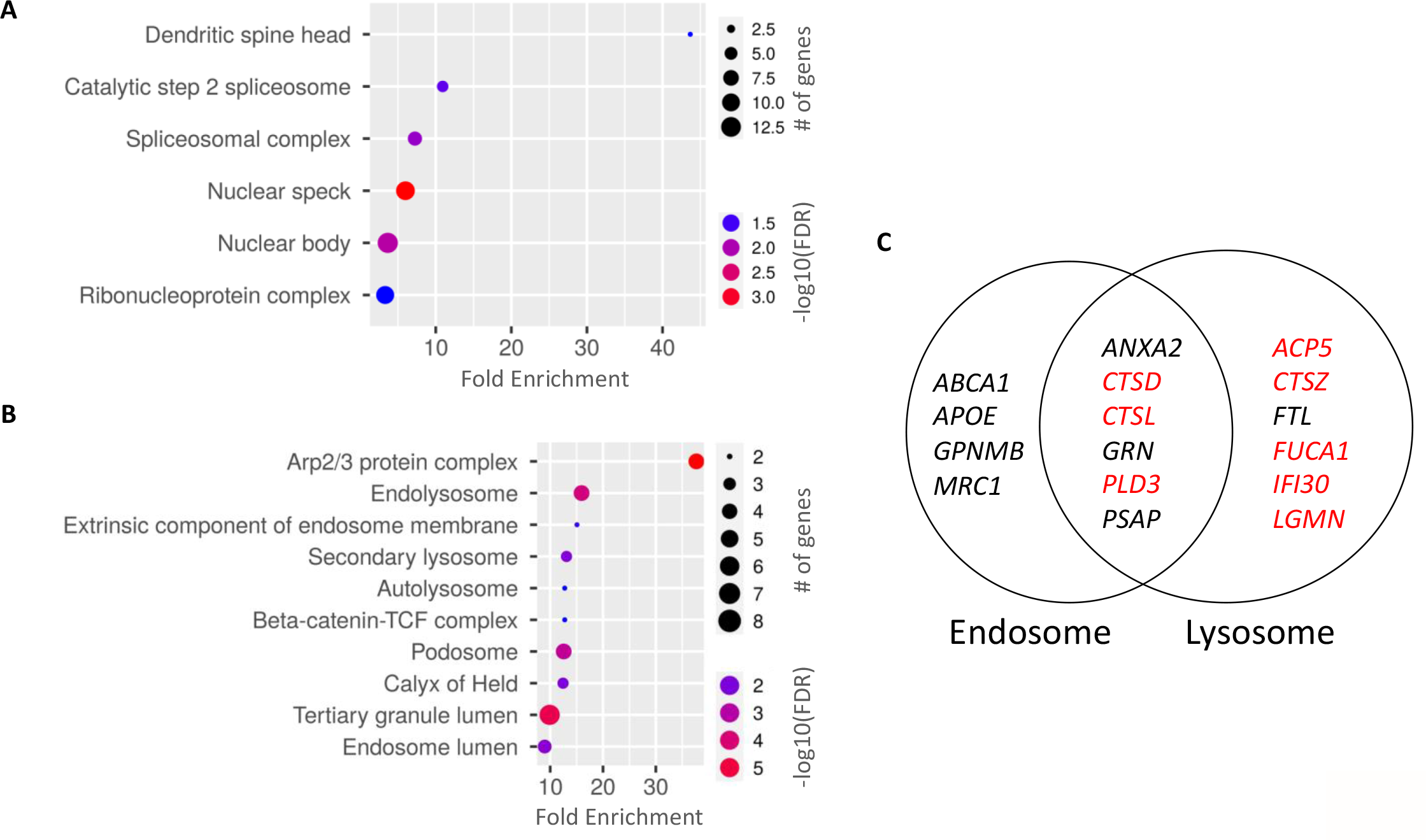
Marker genes for myeloid cells associated with poor GBM patient survival were enriched for cellular components of lysosomes/endolysosomes. **(A)** GO-Cellular Component enrichment analysis of 107 marker genes from MC2 and MC7, the two myeloid cell clusters identified to be associated with better prognosis in glioma patients. Six Cellular Components were identified as enriched (FDR cutoff = 0.05, sorted by fold enrichment). **(B)** GO-Cellular Component enrichment analysis of 309 marker genes from myeloid MC3 and MC5 clusters, the two macrophage clusters identified to be associated with worse prognosis in glioma patients. The chart plot shows the top ten enriched cellular components (FDR cutoff = 0.05, sorted by fold enrichment). (A-B) The size of each dot corresponds to the fold enrichment for each cellular component as shown on the x-axis. **(C)** Illustration of the 16 MC3 marker genes that encode proteins determined to be localized at endosomes and/or lysosomes with high confidence, as denoted by the COMPARTMENTS database (https://compartments.jensenlab.org/Search).

**Fig. 4.**
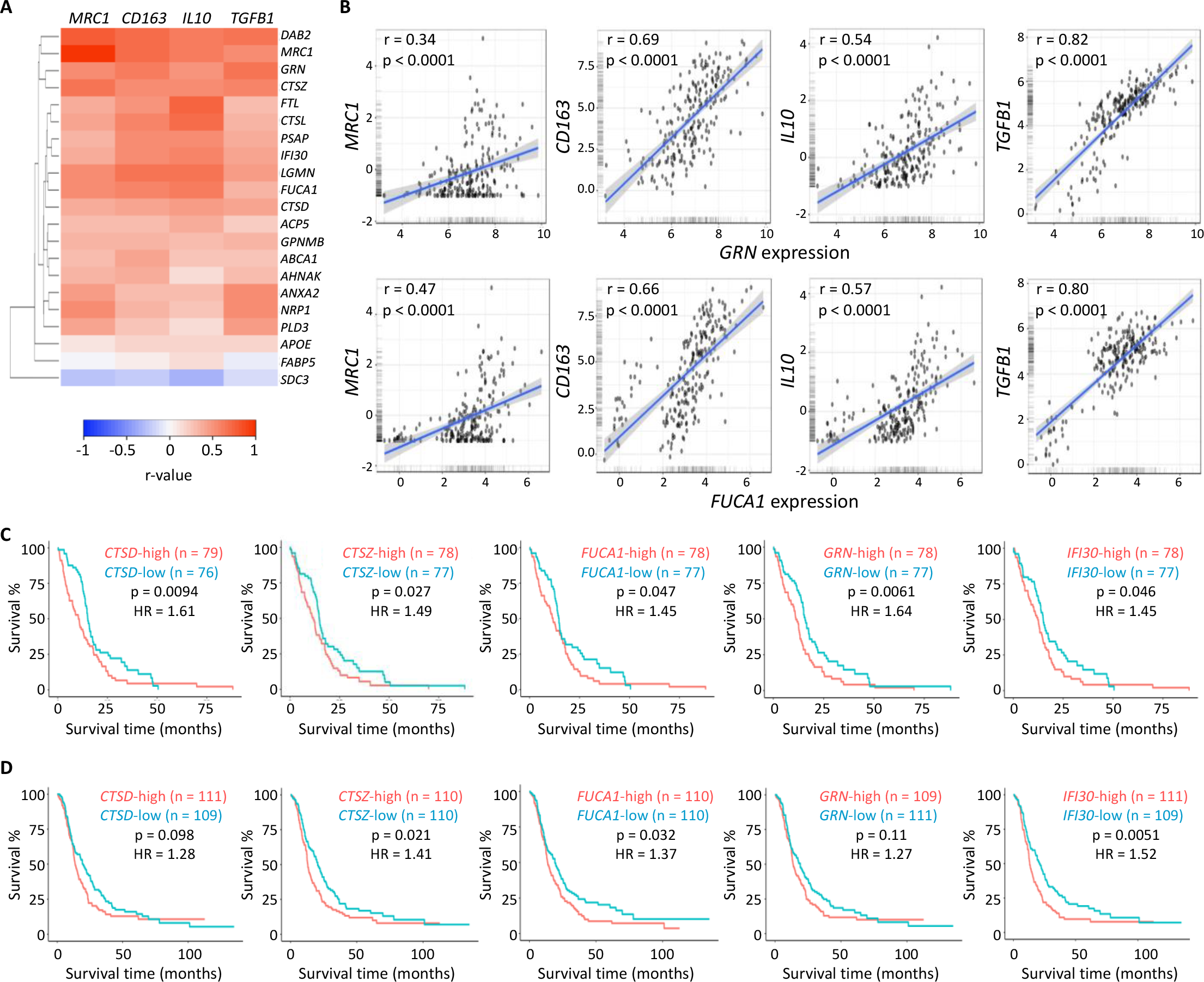
Endolysosomal machinery genes correlate with marker genes indicative of immunosuppression and are associated with worse patient survival in GBMs. **(A)** Heatmap demonstrating the Pearson correlation coefficient values (r-values) between the expression of 21 endolysosomal genes and the immunosuppression markers *MRC1* (CD206), *CD163, IL10*, and *TGFB1*. The heatmap was generated using Heatmapper (clustering method: single linkage; distance measurement method: Euclidean). The TCGA GBM mRNA-seq dataset was used for the correlation analyses (n = 162). **(B)** Scatter plots showing examples of positive correlations between the expression of endolyosomal genes and immunosuppression markers. The CGGA primary GBM dataset was used for the analyses (n = 223). Plots were generated using the GlioVis portal, and r-values and p-values were calculated based on Pearson correlation analysis by the Gliovis portal. **(C)** Kaplan-Meier survival analyses indicating significant correlations between the expression of five genes and GBM patients’ overall survival. The TCGA GBM mRNA-seq dataset was used for the analyses (n = 155). **(D)** Kaplan-Meier survival analyses indicating significant correlations between the expression of five genes and GBM patients’ overall survival. The CGGA primary GBM dataset was used for the analyses (n = 220). (C-D) Median expression was used as the cutoff value for high versus low gene expression. Plots were generated using the GlioVis portal, and p-values and hazard ratios were calculated based on log-rank tests and Cox proportional hazards model, respectively, by the Gliovis portal.

While the findings above suggest the roles of endolysosomes in GBM pathogenesis, particularly in shaping GBM’s immunosuppressive microenvironment, they raise the question of the roles of endolysosomes in the context of GBM resistance to immunotherapy. To address this question, we took advantage of the recent findings that subsets of *SIGLEC9*^+^ TAMs dampen GBM’s response to immunotherapy^15,25^. We specifically focused on two subsets of *SIGLEC9*^+^ TAMs that were found to be highly plastic and immunosuppressive (*SIGLEC9*^+^*MARCO*^+^ TAMs, cluster C9, and *SIGLEC9*^+^*SEPP1*^+^ TAMs, cluster C2)^15^, and examined genes that were upregulated in these two populations of TAMs (versus their respective *SIGLEC9*^-^ counterparts). Enrichment analysis of GO-Cellular Components revealed that in each case, lysosomes/endosomes were enriched only in genes that were upregulated in *SIGLEC9*^+^ TAMs, but not those in their *SIGLEC9*^-^ counterparts (**Supplementary Fig. 5A-B**). Independently, KEGG pathway analysis of genes that defined the *SIGLEC9*^+^ TAMs confirmed lysosomes as the most prominent feature of these immunosuppressive TAMs (**Supplementary Fig. 5C**). Of note, the set of five genes described above (*CTSD, CTSZ, FUCA1, GRN*, and *IFI30*) were among those that were significantly elevated in the immunosuppressive *SIGLEC9*^+^ TAMs^15^. Together with previous findings^15,25^, the results from these analyses suggest that endolysosomes likely play instrumental roles in conferring GBM resistance to immunotherapy.

## Discussion

In summary, by analyzing gene expression profiles, cell type deconvolution, and cellular component enrichment in subsets of non-tumor cells in GBMs, we postulate that the lysosomal machinery is a major component associated with immunosuppressive (pro-tumorigenic) TAMs, and that in addition to their well-documented roles in tumor cells, lysosomes also promote GBM pathogenesis and resistance to immunotherapy through a non-tumor cell-autonomous manner. While these findings were primarily based on suggestive results from gene expression and correlative analyses, they are coherent with prior independent studies. For instance, chloroquine, through disrupting the function of lysosomes, stimulates anti-tumor immunity in a melanoma tumor model by switching M2 macrophages toward M1 phenotypes^26^. Although the microenvironments in GBM and melanoma are expected to be substantially different, the chloroquine-driven M2 to M1 switch provides direct evidence to support the notion that functionally opposite types of macrophages indeed rely differently on the endolysosomal function. Additionally, it is noted that neurodegenerative conditions in humans caused by loss-of-function mutations of genes in the endolysosomal machinery commonly feature microglia activation (i.e., neuroinflammation)^27,28^. The most interesting examples include *GRN* and *FUCA1*, as their loss of functions directly causes lysosomal dysregulation and neuroinflammation, accompanied by neurodegenerative disorders such as Alzheimer’s disease (AD) and frontotemporal dementia (FTD)^29-31^. Separately, *GRN* has been known to be an immunosuppressive protein^32^, and a recent study has linked *FUCA1* to autophagy and macrophage infiltration in gliomagenesis^33^.

Findings presented in this study also inform our understanding of TAMs and have direct implications for targeting the endolysosomal system for cancer treatments in several ways. First, while the heterogeneity and plasticity of TAMs, as defined by their distinct gene expression profiles / featured molecular pathways (e.g., Hallmark pathways), have been well documented in GBMs, they can also be viewed in a highly simplified way as two functionally opposite types (i.e., immunosuppressive/pro-tumorigenic and immunoactive/anti-tumorigenic) that produce different cytokines and/or respond to cytokines in different ways^18,21^. Our cellular component-focused findings suggest that heterogenous and functionally opposite TAMs also depend on distinct cellular organelles for exerting their pro- or anti-tumorigenic effects. This suggests that although lysosomal hydrolysis has been known to be a cellular process critical to macrophages, it is possible that therapeutic targeting of lysosomes can affect immunosuppressive TAMs more than immunoactive TAMs, and shift the TAMs and the GBM microenvironment toward a state favorable for anti-tumor response and immunotherapy. Second, multiple strategies for targeting lysosomes have been devised, including altering the acidic microenvironment (e.g., chloroquine), damaging lysosomal membrane (e.g., cationic amphiphilic drugs), and direct inhibition of hydrolases^6^. In this context, progranulin is particularly interesting. While progranulin is a secreted protein, it can be internalized into cells through cognate receptors and processed into the functional form, termed granulin, in endolysosomes to support the hydrolysis function of these organelles^34-36^. Most importantly, recent studies have revealed that the major mechanistic role of granulin in endolysosomes is to maintain the homeostasis of an unusual, endolysosome-specific family of lipids, bis(monoacylglycero)phosphate (BMP), in these organelles^23,24^. Thus, multiple proteins in this progranulin-BMP axis can be explored for therapeutically targeting the progranulin pathway. Of note, a previous study also suggests that progranulin promotes GBM cell stemness and their resistance to temozolomide, raising the intriguing possibility that targeting the progranulin pathway can provide dual benefits: attenuation of both GBM stemness and immunosuppression. The potential immunosuppressive effect can be tested using *Grn*-knockout mice, which exhibit signs of neuroinflammation but are otherwise viable^37^, as the host mice for tumor generation in future studies. Finally, we propose that, while the relationship between endolysosomes and the TAMs / immune microenvironment in GBMs awaits further investigation, it is warranted to incorporate assays of immune cells and immune microenvironment when assessing endolysosome-based therapeutic approaches in translational research and clinical trials (e.g., chloroquine^38^) for GBM patients.

## Acknowledgement

This work was supported by the Preston Robert Tisch Brain Tumor Center and the Department of Pathology at Duke University, the National Institute of Neurological Disorders and Stroke (NINDS) at the National Institutes of Health (NIH) (NS101074), and a pilot research grant from Duke Cancer Institute as part of the NIH National Cancer Institute P30 Cancer Center Support Grant (Grant ID: NIH CA014236).

## References

1 Shin, H. R. & Zoncu, R. The Lysosome at the Intersection of Cellular Growth and Destruction. Dev Cell 54, 226–238, doi:10.1016/j.devcel.2020.06.010 (2020).

2 Bright, N. A., Davis, L. J. & Luzio, J. P. Endolysosomes Are the Principal Intracellular Sites of Acid Hydrolase Activity. Current biology : CB 26, 2233–2245, doi:10.1016/j.cub.2016.06.046 (2016).

3 Iulianna, T., Kuldeep, N. & Eric, F. The Achilles’ heel of cancer: targeting tumors via lysosome-induced immunogenic cell death. Cell Death Dis 13, 509, doi:10.1038/s41419-022-04912-8 (2022).

4 Machado, E. R., Annunziata, I., van de Vlekkert, D., Grosveld, G. C. & d’Azzo, A. Lysosomes and Cancer Progression: A Malignant Liaison. Front Cell Dev Biol 9, 642494, doi:10.3389/fcell.2021.642494 (2021).

5 Rebecca, V. W. et al. A Unified Approach to Targeting the Lysosome’s Degradative and Growth Signaling Roles. Cancer discovery 7, 1266–1283, doi:10.1158/2159-8290.CD-17-0741 (2017).

6 Piao, S. & Amaravadi, R. K. Targeting the lysosome in cancer. Ann N Y Acad Sci 1371, 45–54, doi:10.1111/nyas.12953 (2016).

7 Racle, J. & Gfeller, D. EPIC: A Tool to Estimate the Proportions of Different Cell Types from Bulk Gene Expression Data. Methods in molecular biology 2120, 233–248, doi:10.1007/978-1-0716-0327-7_17 (2020).

8 Finotello, F. et al. Molecular and pharmacological modulators of the tumor immune contexture revealed by deconvolution of RNA-seq data. Genome Med 11, 34, doi:10.1186/s13073-019-0638-6 (2019).

9 Aran, D., Hu, Z. & Butte, A. J. xCell: digitally portraying the tissue cellular heterogeneity landscape. Genome biology 18, 220, doi:10.1186/s13059-017-1349-1 (2017).

10 Newman, A. M. et al. Robust enumeration of cell subsets from tissue expression profiles. Nature methods 12, 453–457, doi:10.1038/nmeth.3337 (2015).

11 Li, T. et al. TIMER2.0 for analysis of tumor-infiltrating immune cells. Nucleic acids research 48, W509–W514, doi:10.1093/nar/gkaa407 (2020).

12 Ge, S. X., Jung, D. & Yao, R. ShinyGO: a graphical gene-set enrichment tool for animals and plants. Bioinformatics 36, 2628–2629, doi:10.1093/bioinformatics/btz931 (2020).

13 Raudvere, U. et al. g:Profiler: a web server for functional enrichment analysis and conversions of gene lists (2019 update). Nucleic acids research 47, W191–W198, doi:10.1093/nar/gkz369 (2019).

14 Bowman, R. L., Wang, Q., Carro, A., Verhaak, R. G. & Squatrito, M. GlioVis data portal for visualization and analysis of brain tumor expression datasets. Neuro-oncology 19, 139–141, doi:10.1093/neuonc/now247 (2017).

15 Mei, Y. et al. Siglec-9 acts as an immune-checkpoint molecule on macrophages in glioblastoma, restricting T-cell priming and immunotherapy response. Nat Cancer, doi:10.1038/s43018-023-00598-9 (2023).

16 Zhao, Z. et al. Chinese Glioma Genome Atlas (CGGA): A Comprehensive Resource with Functional Genomic Data from Chinese Glioma Patients. Genomics Proteomics Bioinformatics 19, 1–12, doi:10.1016/j.gpb.2020.10.005 (2021).

17 Babicki, S. et al. Heatmapper: web-enabled heat mapping for all. Nucleic acids research 44, W147–153, doi:10.1093/nar/gkw419 (2016).

18 Khan, F. et al. Macrophages and microglia in glioblastoma: heterogeneity, plasticity, and therapy. The Journal of clinical investigation 133, doi:10.1172/JCI163446 (2023).

19 Buonfiglioli, A. & Hambardzumyan, D. Macrophages and microglia: the cerberus of glioblastoma. Acta Neuropathol Commun 9, 54, doi:10.1186/s40478-021-01156-z (2021).

20 Dapash, M., Hou, D., Castro, B., Lee-Chang, C. & Lesniak, M. S. The Interplay between Glioblastoma and Its Microenvironment. Cells 10, doi:10.3390/cells10092257 (2021).

21 Abdelfattah, N. et al. Single-cell analysis of human glioma and immune cells identifies S100A4 as an immunotherapy target. Nature communications 13, 767, doi:10.1038/s41467-022-28372-y (2022).

22 Binder, J. X. et al. COMPARTMENTS: unification and visualization of protein subcellular localization evidence. Database (Oxford) 2014, bau012, doi:10.1093/database/bau012 (2014).

23 Logan, T. et al. Rescue of a lysosomal storage disorder caused by Grn loss of function with a brain penetrant progranulin biologic. Cell 184, 4651–4668 e4625, doi:10.1016/j.cell.2021.08.002 (2021).

24 Boland, S. et al. Deficiency of the frontotemporal dementia gene GRN results in gangliosidosis. Nature communications 13, 5924, doi:10.1038/s41467-022-33500-9 (2022).

25 Schmassmann, P. et al. Targeting the Siglec-sialic acid axis promotes antitumor immune responses in preclinical models of glioblastoma. Science translational medicine 15, eadf5302, doi:10.1126/scitranslmed.adf5302 (2023).

26 Chen, D. et al. Chloroquine modulates antitumor immune response by resetting tumor-associated macrophages toward M1 phenotype. Nature communications 9, 873, doi:10.1038/s41467-018-03225-9 (2018).

27 Stephenson, J., Nutma, E., van der Valk, P. & Amor, S. Inflammation in CNS neurodegenerative diseases. Immunology 154, 204–219, doi:10.1111/imm.12922 (2018).

28 Guzman-Martinez, L. et al. Neuroinflammation as a Common Feature of Neurodegenerative Disorders. Front Pharmacol 10, 1008, doi:10.3389/fphar.2019.01008 (2019).

29 Stepien, K. M., Ciara, E. & Jezela-Stanek, A. Fucosidosis-Clinical Manifestation, Long-Term Outcomes, and Genetic Profile-Review and Case Series. Genes (Basel) 11, doi:10.3390/genes11111383 (2020).

30 Wolf, H. et al. A mouse model for fucosidosis recapitulates storage pathology and neurological features of the milder form of the human disease. Dis Model Mech 9, 1015–1028, doi:10.1242/dmm.025122 (2016).

31 Nalls, M. A. et al. Evidence for GRN connecting multiple neurodegenerative diseases. Brain Commun 3, fcab095, doi:10.1093/braincomms/fcab095 (2021).

32 Jian, J., Konopka, J. & Liu, C. Insights into the role of progranulin in immunity, infection, and inflammation. Journal of leukocyte biology 93, 199–208, doi:10.1189/jlb.0812429 (2013).

33 Xu, L. et al. Downregulation of alpha-l-fucosidase 1 suppresses glioma progression by enhancing autophagy and inhibiting macrophage infiltration. Cancer science 111, 2284–2296, doi:10.1111/cas.14427 (2020).

34 Hu, F. et al. Sortilin-mediated endocytosis determines levels of the frontotemporal dementia protein, progranulin. Neuron 68, 654–667, doi:10.1016/j.neuron.2010.09.034 (2010).

35 Zhou, X. et al. Lysosomal processing of progranulin. Mol Neurodegener 12, 62, doi:10.1186/s13024-017-0205-9 (2017).

36 Zhou, X. et al. Progranulin deficiency leads to reduced glucocerebrosidase activity. PloS one 14, e0212382, doi:10.1371/journal.pone.0212382 (2019).

37 Yin, F. et al. Exaggerated inflammation, impaired host defense, and neuropathology in progranulin-deficient mice. The Journal of experimental medicine 207, 117–128, doi:10.1084/jem.20091568 (2010).

38 Weyerhauser, P., Kantelhardt, S. R. & Kim, E. L. Re-purposing Chloroquine for Glioblastoma: Potential Merits and Confounding Variables. Front Oncol 8, 335, doi:10.3389/fonc.2018.00335 (2018).

