## Supplementary Figures for "Gene expression analysis suggests immunosuppressive roles of endolysosomes in glioblastoma"

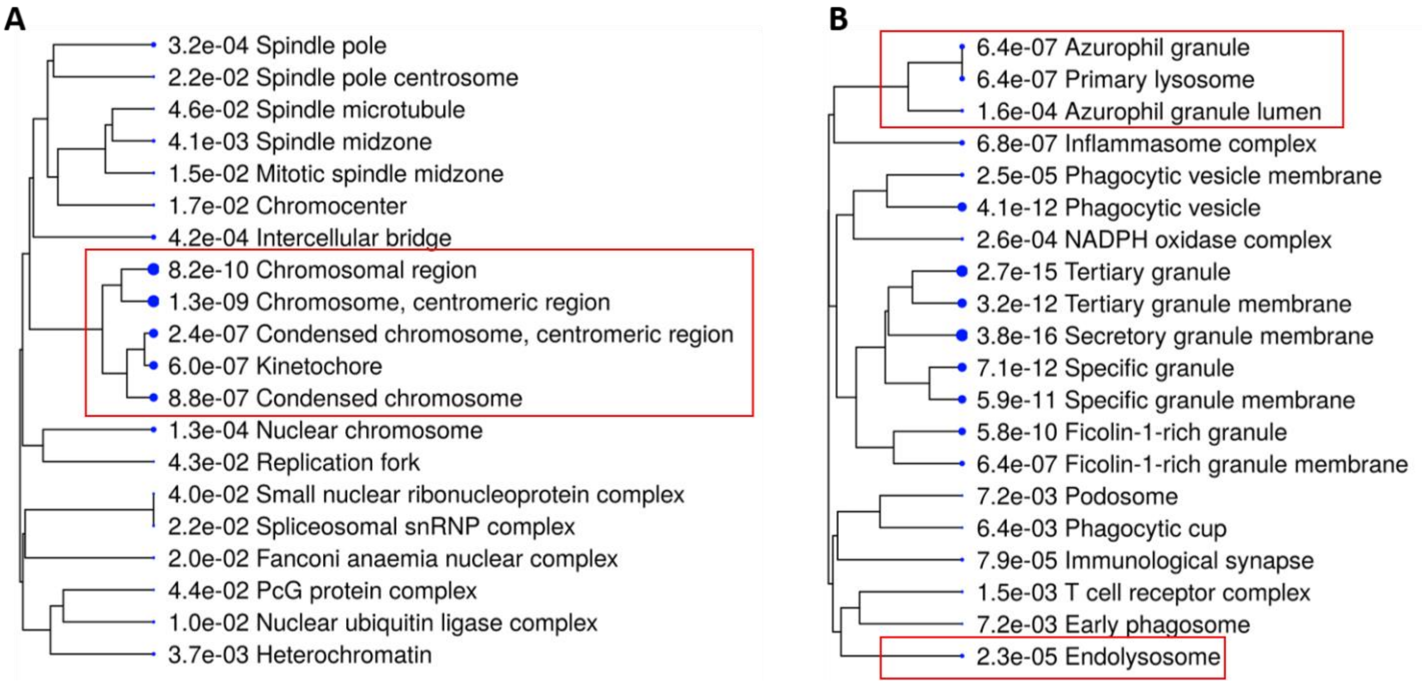


**Suppl Fig. 1. Gene expression analysis suggests tumor and non-tumor cells in GBMs featured distinct cellular components.** The mRNA-seq data (protein-coding genes, tpm) of 162 TCGA GBM samples were subjected to quanTIseq analysis for estimating fractions of each type of cells. **(A)** GO-Cellular Component enrichment analysis of genes positively correlated with cancer cells (annotated as "Other Cells") (r >= 0.3; n = 290). **(B)** GO-Cellular Component enrichment analysis of genes negatively correlated with cancer cells (annotated as "Other Cells") (r <= -0.5; n = 347). (A-B) Dendrograms show the top 20 cellular components sorted by fold enrichment, and the size of each dot denotes the relative number of genes in each cellular component.

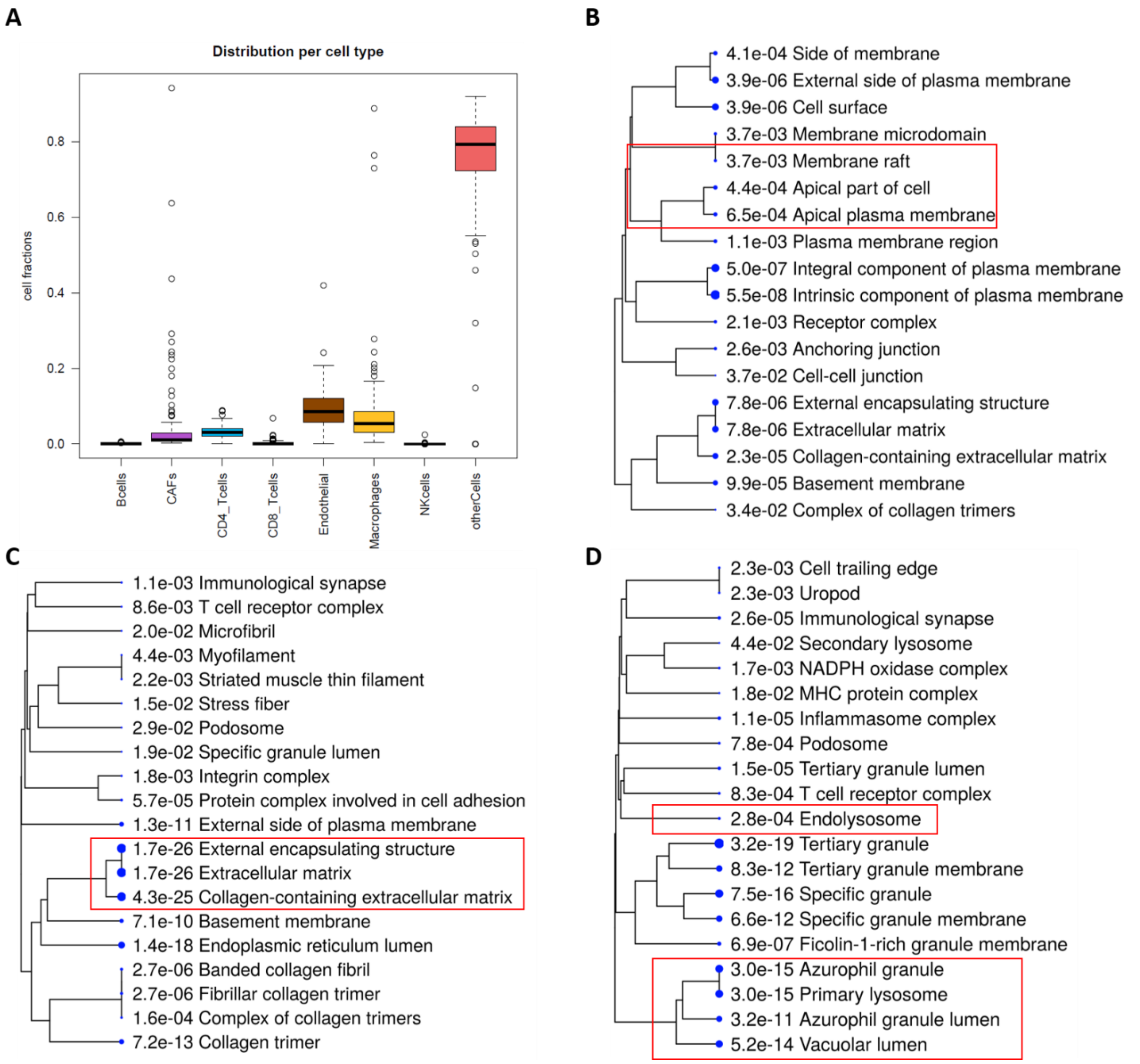


**Suppl. Fig. 2. Tumor-associated macrophages are the major cell type that contributes to the prominent feature of endolysosomal machinery in GBMs. (A-C)** The mRNA-seq data (protein-coding genes, tpm) of 162 TCGA GBM samples were subjected to EPIC analysis for estimating fractions of each type of cells. **(A)** Box plot showing the fractions of different cell types, including infiltrating immune cells and other cells (i.e., cancer cells), as estimated by EPIC. **(B)** GO-Cellular Component enrichment analysis of genes positively correlated with endothelial cells (r >= 0.3; n = 237). **(C)** GO-Cellular Component enrichment analysis of genes positively correlated with CAF (r >= 0.5; n = 543). **(D)** The fractions of macrophages were estimated by xCELL using the mRNA-seq data of 162 TCGA GBM samples, and genes positively correlated with macrophage (r >= 0.5) were used for GO-Cellular Component enrichment analysis. (B-D) Dendrograms show the top 20 cellular components sorted by fold enrichment, and the size of each dot denotes the relative number of genes in each cellular component.

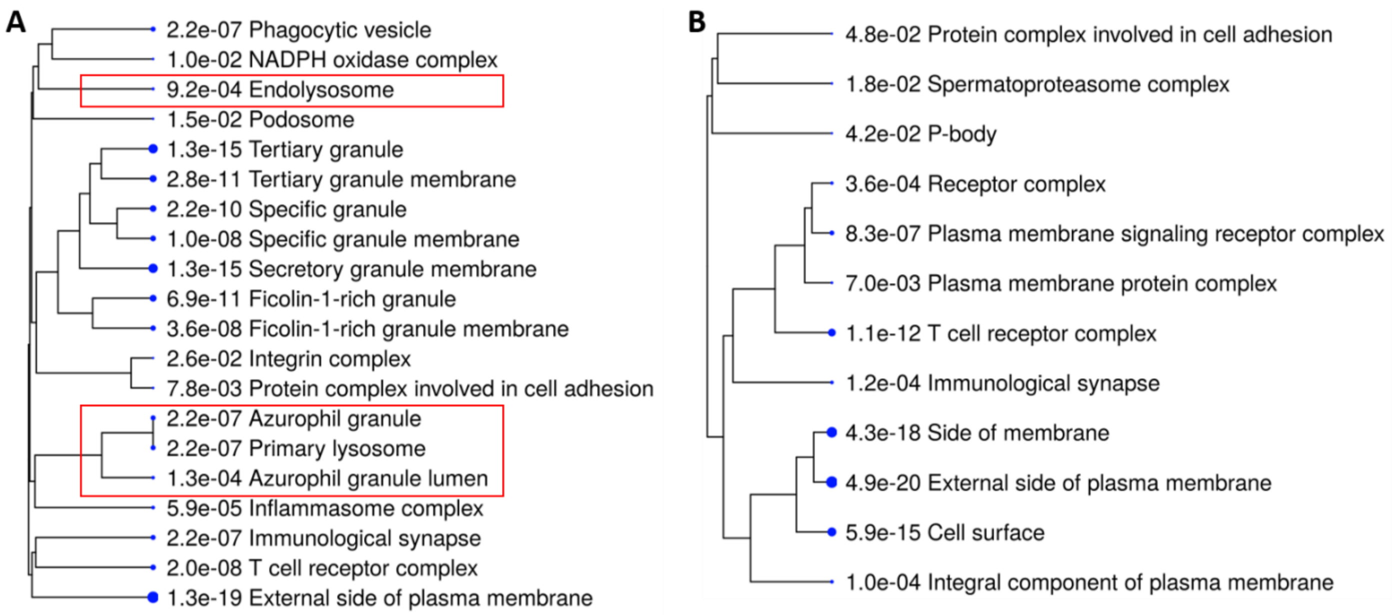


C

| myeloid cluster 5 marker genes (n=254) | | | | myeloid cluster 3 genes (n=55) | | | |
|---|---|---|---|---|---|---|---|
| Enrichment FDR | # of genes | Fold Enrichment | GO-cellular Component | Enrichment FDR | # of genes | Fold Enrichment | GO-cellular Component |
| 1.60E-16 | 50 | 4.3 | Secretory vesicle | 2.40E-10 | 17 | 8.3 | Lytic vacuole |
| 3.80E-18 | 49 | 4.9 | Secretory granule | 2.40E-10 | 17 | 8.3 | Lysosome |
| 2.30E-13 | 40 | 4.3 | Anchoring junction | 1.20E-09 | 17 | 7.4 | Vacuole |
| 6.00E-17 | 34 | 7.1 | Focal adhesion | 1.20E-09 | 17 | 7.4 | Secretory granule |
| 6.30E-17 | 34 | 7 | Cell-substrate junction | 1.40E-08 | 17 | 6.2 | Secretory vesicle |
| 7.50E-05 | 33 | 2.3 | Cytoplasmic vesicle membrane | 9.00E-12 | 15 | 14.2 | Collagen-containing extracellular matrix |
| 9.40E-05 | 33 | 2.3 | Vesicle membrane | 2.40E-10 | 15 | 10.5 | External encapsulating structure |
| 2.50E-12 | 30 | 5.4 | Actin cytoskeleton | 2.40E-10 | 15 | 10.6 | Extracellular matrix |
| 1.40E-06 | 30 | 3 | Vacuole | 3.50E-05 | 14 | 4.3 | Cytoplasmic vesicle membrane |
| 1.40E-06 | 28 | 3.2 | Lytic vacuole | 3.80E-05 | 14 | 4.2 | Vesicle membrane |
| 1.40E-06 | 28 | 3.2 | Lysosome | 3.10E-13 | 13 | 26.8 | Vacuolar lumen |
| 1.00E-13 | 27 | 7.2 | Secretory granule lumen | 3.40E-05 | 13 | 4.7 | Endosome |
| 1.10E-13 | 27 | 7.1 | Vesicle lumen | 9.20E-05 | 13 | 4.1 | Plasma membrane region |
| 1.10E-13 | 27 | 7.1 | Cytoplasmic vesicle lumen | 2.90E-05 | 12 | 5.3 | Cell surface |
| 3.80E-07 | 24 | 3.9 | External encapsulating structure | 1.70E-08 | 11 | 12.6 | Secretory granule lumen |
| 3.80E-07 | 24 | 3.9 | Extracellular matrix | 1.70E-08 | 11 | 12.4 | Vesicle lumen |
| 4.30E-02 | 23 | 1.7 | Plasma membrane region | 1.70E-08 | 11 | 12.5 | Cytoplasmic vesicle lumen |
| 4.60E-08 | 22 | 4.9 | Cell leading edge | 2.40E-10 | 9 | 33.3 | Lysosomal lumen |
| 4.60E-08 | 22 | 4.8 | Collagen-containing extracellular matrix | 1.80E-04 | 7 | 8.2 | Membrane raft |
| 2.90E-03 | 22 | 2.3 | Cell surface | 1.80E-04 | 7 | 8.2 | Membrane microdomain |

**Suppl. Fig. 3. Different subsets of myeloid cells featured distinct cellular components. (A-B)** The mRNA-seq data (protein-coding genes, tpm) of 162 TCGA GBM samples were subjected to CIBERSORT (absolute mode) analysis for estimating fractions of M1 and M2 macrophages. **(A)** GO-Cellular Component enrichment analysis of genes positively correlated with M2 macrophages (r >= 0.5; n = 429). **(B)** GO-Cellular Component enrichment analysis of genes positively correlated with M1 macrophages (r >= 0.3; n = 318). Note that the cutoff value of r >= 0.3 was used for the M1 macrophage analysis to have the number of analyzed genes more comparable to the M2 macrophage analysis (when the cutoff value of r >= 0.5 was used, only 56 genes were identified). (A-B) Dendrograms show the significantly enriched cellular components (FDR cutoff = 0.05), and the size of each dot denotes the relative number of genes in each cellular component. **(C)** GO-Cellular Component enrichment analysis of marker genes for MC3 or MC5. The table shows the top 20 cellular components sorted by the number of genes identified. Cellular components directly indicative of lysosomes/endosomes were highlighted in red.

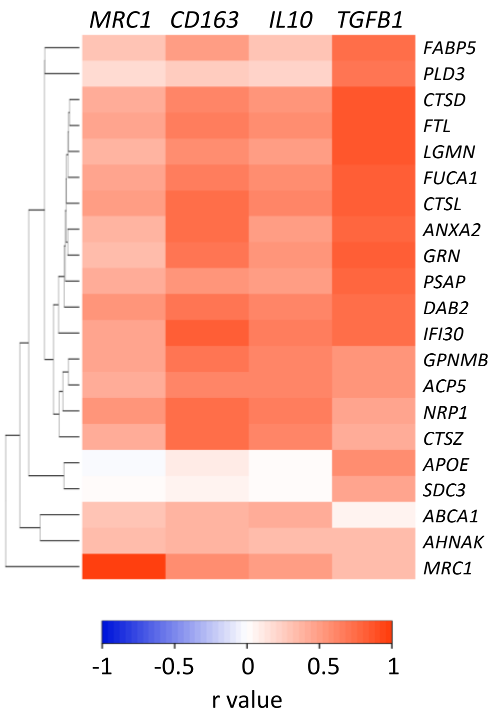


**Suppl. Fig. 4. Endolysosomal machinery genes correlate with marker genes indicative of immunosuppression.** Heatmap demonstrating the Pearson correlation coefficient values (r-values) between the expression of 21 endolysosomal genes and immunosuppression markers *MRC1* (CD206), *CD163*, *IL10,* and *TGFB1*. The heatmap was generated using Heatmapper (clustering method: single linkage; distance measurement method: Euclidean). The CGGA primary GBM dataset was used for the correlation analysis (n = 223).

**A**

| genes upregulated in SIGLEC9$^+$MARCO$^+$ TAMs (n=374) | | | |
|---|---|---|---|
| Enrichment FDR | # of genes | Fold Enrichment | GO-cellular component |
| 3.10E-34 | 86 | 5 | Secretory vesicle |
| 1.20E-37 | 84 | 5.8 | Secretory granule |
| 3.30E-14 | 64 | 3.1 | Vesicle membrane |
| 5.70E-14 | 63 | 3 | Cytoplasmic vesicle membrane |
| 1.50E-18 | 57 | 4.2 | Anchoring junction |
| 6.80E-05 | 54 | 1.8 | Intrinsic component of plasma membrane |
| 1.70E-04 | 51 | 1.8 | Integral component of plasma membrane |
| 2.20E-11 | 47 | 3.3 | Cell surface |
| 9.10E-04 | 47 | 1.7 | Mitochondrion |
| 1.10E-21 | 46 | 6.5 | Cell-substrate junction |
| 3.40E-21 | 45 | 6.4 | Focal adhesion |
| 3.10E-10 | 45 | 3.1 | Vacuole |
| 1.10E-10 | 43 | 3.3 | Lytic vacuole |
| 1.10E-10 | 43 | 3.3 | Lysosome |
| 1.20E-05 | 43 | 2.2 | Plasma membrane region |
| 1.30E-04 | 42 | 2 | Membrane protein complex |
| 2.00E-04 | 42 | 2 | Neuron projection |
| 7.60E-21 | 40 | 7.3 | Secretory granule lumen |
| 9.50E-21 | 40 | 7.2 | Cytoplasmic vesicle lumen |
| 9.90E-21 | 40 | 7.2 | Vesicle lumen |

| genes upregulated in SIGLEC9$^-$MARCO$^-$ TAMs (n=174) | | | |
|---|---|---|---|
| Enrichment FDR | # of genes | Fold Enrichment | GO-cellular component |
| 1.20E-03 | 25 | 2.7 | Neuron projection |
| 1.20E-03 | 25 | 2.7 | Synapse |
| 7.90E-03 | 21 | 2.2 | Supramolecular complex |
| 5.00E-02 | 21 | 1.8 | Golgi apparatus |
| 3.70E-03 | 19 | 2.7 | Supramolecular fiber |
| 3.80E-03 | 19 | 2.6 | Supramolecular polymer |
| 2.10E-02 | 18 | 2.1 | Chromatin |
| 3.10E-02 | 18 | 2 | Nuclear protein-containing complex |
| 3.70E-03 | 17 | 2.9 | Somatodendritic compartment |
| 1.20E-03 | 16 | 3.6 | Dendritic tree |
| 1.20E-03 | 15 | 3.8 | Cell body |
| 2.20E-03 | 15 | 3.4 | Dendrite |
| 2.20E-03 | 13 | 3.7 | Neuronal cell body |
| 8.30E-03 | 13 | 2.9 | Postsynapse |
| 8.80E-03 | 13 | 2.9 | Axon |
| 5.50E-03 | 12 | 3.4 | Actin cytoskeleton |
| 5.50E-03 | 11 | 3.6 | Transcription regulator complex |
| 1.90E-02 | 11 | 2.8 | External encapsulating structure |
| 1.90E-02 | 11 | 2.8 | Extracellular matrix |
| 9.30E-03 | 10 | 3.5 | Cell leading edge |

**B**

| genes upregulated in SIGLEC9$^+$SEPP1$^+$ TAMs (n=445) | | | |
|---|---|---|---|
| Enrichment FDR | # of genes | Fold Enrichment | GO-cellular component |
| 1.80E-08 | 68 | 2.2 | Intrinsic component of plasma membrane |
| 6.30E-22 | 67 | 4.4 | Secretory granule |
| 2.80E-18 | 67 | 3.7 | Secretory vesicle |
| 9.00E-14 | 66 | 3 | Vesicle membrane |
| 3.60E-13 | 64 | 2.9 | Cytoplasmic vesicle membrane |
| 1.90E-14 | 55 | 3.6 | Vacuole |
| 3.10E-15 | 53 | 3.9 | Lytic vacuole |
| 3.10E-15 | 53 | 3.9 | Lysosome |
| 6.50E-11 | 48 | 3.2 | Cell surface |
| 3.60E-08 | 48 | 2.6 | Endosome |
| 3.80E-03 | 47 | 1.6 | Golgi apparatus |
| 1.60E-05 | 45 | 2.2 | Plasma membrane region |
| 3.40E-04 | 43 | 1.9 | Membrane protein complex |
| 1.10E-07 | 40 | 2.8 | Anchoring junction |
| 2.60E-08 | 35 | 3.3 | Side of membrane |
| 7.60E-13 | 31 | 5.5 | Secretory granule membrane |
| 2.40E-07 | 30 | 3.3 | Vacuolar membrane |
| 5.90E-11 | 29 | 5 | Secretory granule lumen |
| 6.70E-11 | 29 | 4.9 | Cytoplasmic vesicle lumen |
| 7.20E-11 | 29 | 4.9 | Vesicle lumen |

| genes upregulated in SIGLEC9$^-$SEPP1$^-$ TAMs (n=266) | | | |
|---|---|---|---|
| Enrichment FDR | # of genes | Fold Enrichment | GO-cellular component |
| 6.90E-11 | 43 | 3.4 | Organelle envelope |
| 6.90E-11 | 43 | 3.4 | Envelope |
| 1.20E-06 | 42 | 2.5 | Mitochondrion |
| 2.40E-11 | 35 | 4.3 | Mitochondrial envelope |
| 8.20E-11 | 33 | 4.3 | Mitochondrial membrane |
| 1.80E-09 | 27 | 4.6 | Organelle inner membrane |
| 9.40E-03 | 26 | 1.9 | Membrane protein complex |
| 4.00E-09 | 25 | 4.8 | Mitochondrial inner membrane |
| 4.60E-02 | 25 | 1.7 | Catalytic complex |
| 2.60E-02 | 24 | 1.8 | Nuclear protein-containing complex |
| 4.10E-02 | 24 | 1.8 | Synapse |
| 5.90E-03 | 22 | 2.2 | Nucleolus |
| 6.90E-11 | 21 | 7.3 | Mitochondrial protein-containing complex |
| 1.20E-14 | 20 | 12.6 | Inner mitochondrial membrane protein complex |
| 1.10E-03 | 20 | 2.7 | Ribonucleoprotein complex |
| 9.70E-03 | 20 | 2.2 | Secretory granule |
| 1.00E-04 | 19 | 3.4 | External encapsulating structure |
| 1.00E-04 | 19 | 3.4 | Extracellular matrix |
| 6.70E-06 | 18 | 4.3 | Collagen-containing extracellular matrix |
| 2.50E-25 | 16 | 73.8 | Mitochondrial proton-transporting ATP synthase complex |

**C**

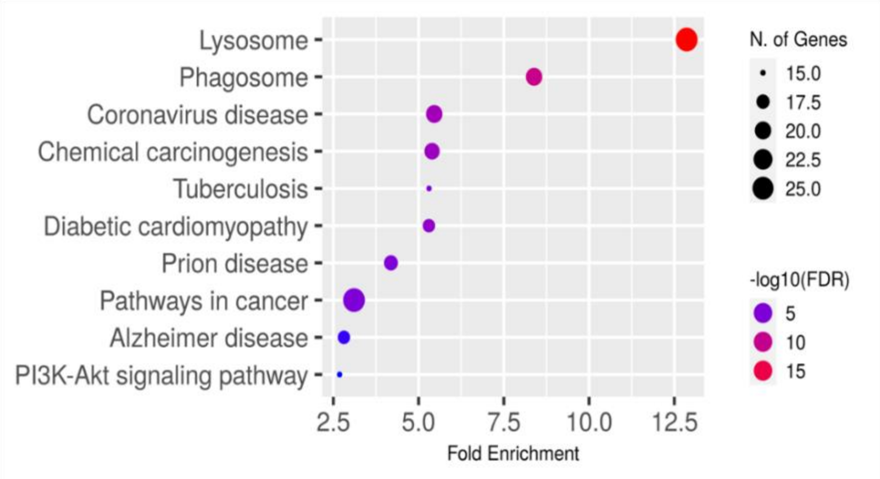


**Suppl. Fig. 5. Lysosomes/endosomes were featured in *SIGLEC9*$^+$ AMs but not in their *SIGLEC9*$^-$ counterparts. (A)** GO-Cellular Component enrichment analysis of genes upregulated in

*SIGLEC9*$^{+}$*MARCO*$^{+}$ TAMs (left panel) or *SIGLEC9*$^{-}$*MARCO*$^{-}$ TAMs (right panel). **(B)** GO-Cellular Component enrichment analysis of genes upregulated in *SIGLEC9*$^{+}$*SEPP1*$^{+}$ TAMs (left panel) or *SIGLEC9*$^{-}$*SEPP1*$^{-}$ TAMs (right panel). (A-B) Dendrograms show the top 20 cellular components sorted by the number of genes identified. Cellular components directly indicative of lysosomes/endosomes were highlighted in red. **(C)** KEGG pathway analysis of genes upregulated in *SIGLEC9*$^{+}$*SEPP1*$^{+}$ TAMs. The chart plot shows the top 10 enriched pathways (FDR cutoff = 0.05, sorted by fold enrichment). Similar analysis of genes upregulated in *SIGLEC9*$^{+}$*MARCO*$^{+}$ TAMs also identified lysosomes as significantly enriched. All known protein-coding genes were used as the background gene set. (A-C) For identifying upregulated genes, an FDR cutoff value of 0.05 was used for all analyses.
